# Memory reactivation during sleep promotes structure abstraction

**DOI:** 10.64898/2026.04.10.717748

**Authors:** Sarah H. Solomon, Sirisha Krishnamurthy, Elizabeth M. Siefert, Claudia M. Gonciulea, Anna C. Schapiro

## Abstract

We readily detect structure in our environments, which in turn guides future learning. Disentangling this structure from the superficial features of a specific learning environment provides an especially strong basis for future generalization, but it remains unclear when and how this kind of abstraction occurs. Memory reactivation during sleep has been hypothesized to support such abstraction, but this has yet to be directly tested. Here we examined this hypothesis by teaching participants novel categories in which patterns of feature covariation were governed by different graph structures. Participants then learned a new category, defined by entirely different features, whose structure was either congruent or incongruent with a previously learned category. If structural knowledge is abstracted away from superficial features, it should facilitate transfer when structures are congruent. In Experiment 1, when two categories were learned in immediate succession, participants showed no transfer benefit, suggesting that structure understanding remained tied to the original features. In Experiment 2, we tested whether offline processing promotes abstraction. Participants either remained awake between learning phases spaced 3 hours apart, or took a nap during which a previously learned category was reactivated using targeted memory reactivation (TMR). Transfer benefits emerged only when the reactivated and target categories shared the same structure, and these benefits increased with the number of cues presented during slow-wave sleep. These findings provide the first direct evidence that memory reactivation during sleep promotes the abstraction of structure, enabling knowledge to transfer across learning episodes with no overlap in features.

## INTRODUCTION

Human intelligence enables us to extract patterns that go beyond surface details, allowing us to generalize to new situations. Whether we are learning semantic categories, navigating environments, or mapping social relationships, success often involves uncovering the deeper structures that organize our experiences. When, why, and how such structures are uncovered remains a central question in cognitive science. Classic accounts of scientific discovery, such as August Kekulé’s report that the ring structure of benzene emerged in a dream, have long suggested that structural insight may not arise during initial learning, but may instead emerge later—perhaps especially during sleep.

Sleep helps to mold our recent experiences into long-term memories offline. This consolidation process does more than merely crystallize transient memories into durable ones, with a growing literature suggesting that sleep also promotes memory *transformation* (Landmann et al., 2014; Stickgold & Walker, 2013). Sleep has been linked to phenomena such as insight and rule-extraction (Wagner et al., 2004; Pace-Schott, 2012), relational memory and inference (Ellenbogen et al., 2007; Laur et al., 2010), and statistical learning and transfer (Gomez et al., 2006; Durrant et al., 2011; Albouy et al., 2013; Durrant et al., 2016; Garber & Fiser, 2025). In the semantic learning context, sleep has been found to facilitate probabilistic category learning (Djonlagic et al., 2009), aid semantic learning and generalization (Friedrich et al., 2015; Lau et al., 2011), and strengthen or weaken individual features in a newly learned category (Siefert et al., 2024; Sherman et al., 2025). Thus, after sleep, a category representation is not simply a strengthened version of the original memory, but a representation that has been sculpted and transformed. Despite longstanding assumptions that one consequence of these sleep-dependent transformations is the emergence of abstract structure (as in Kekulé’s benzene “ring” anecdote), the claim that sleep supports structure abstraction has not been directly tested.

Memory replay or reactivation is one potential mechanism for this sleep-based memory transformation (Inostroza & Born, 2013). Targeted memory reactivation (TMR) is an experimental paradigm in which memory-related sensory cues (e.g., sounds, odors) are reintroduced during sleep to promote reactivation of recently learned content. TMR benefits memory for previously learned items (e.g., Rasch et al., 2007; Rudoy et al., 2009; Oudiette & Paller, 2013; Hu et al., 2020) and contributes to the kinds of memory transformations described above (Siefert et al. 2024; Sherman et al., 2025), but its ability to promote structure abstraction in particular has not yet been assessed (Fig. 1).

**Fig 1:**
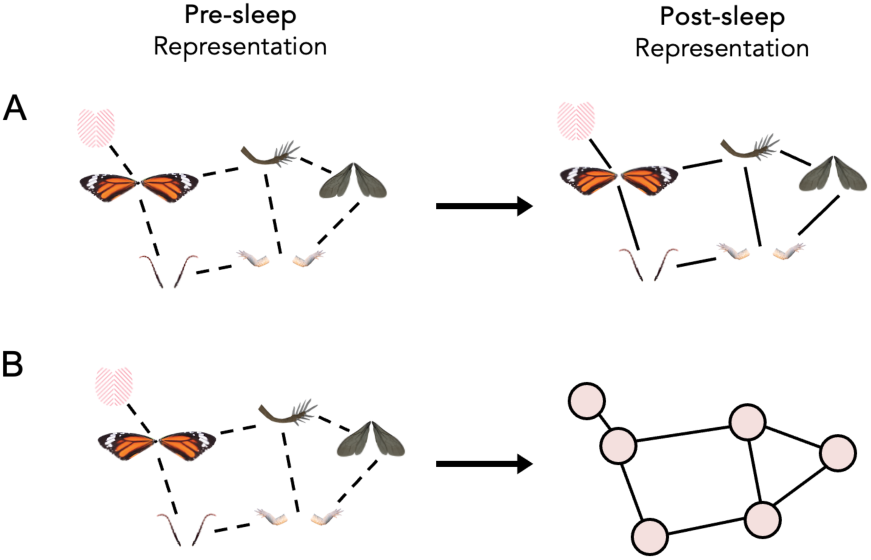
Structure Abstraction: Consolidation processes during wake or sleep could (A) strengthen direct feature relationships and/or (B) disentangle the features from the structure so that this abstracted structure may be used in new learning contexts.

Domains can often be well-described by a certain structural form: living things and kinship systems correspond to trees; colors, seasons, and times of day are rings; and social relationships and academic disciplines are well-described by community structure, in which tightly connected clusters are embedded within a broader network. Semantic structure also exists at a more granular level—the level of semantic features. Features exhibit patterns of association, and this structure supports both our world knowledge and the inferences we can make. For example, bouncy things are often round, metal things are often electrical, and things that have roots often need sunlight. Feature associations can be leveraged to acquire knowledge of structured semantic domains in humans and artificial neural networks, can explain judgments of semantic similarity and typicality, and can also explain the trajectory of semantic decline in age and disease (Rogers & McClelland, 2004). Feature associations also play a role in the acquisition and representation of individual semantic categories. Humans are sensitive to pairwise feature correlations during category learning (Medin et al., 1982; Wattenmaker, 1991; 1993) and are also sensitive to the higher-level structure of feature covariation within a novel category (Solomon & Schapiro, 2024).

Although humans can uncover deep structures, it remains unclear to what extent these structures are represented in an abstract form versus remaining tied to specific items or surface features. On the one hand, exemplar-based theories of learning argue that representations are bound to individual items, with structural knowledge only emerging as a function of the psychological distances between them (Nosofsky, 1986; Medin & Schaffer, 1978); this class of theories does not posit abstract structural representations. On the other hand, some researchers evoke cognitive maps (Tolman, 1948), which specify the relative spatial relations between items in a Euclidean 2D space. Cognitive maps instantiate a form of structural abstraction, particularly when theoretically grounded in the neural grid-like codes in the hippocampal system, since this same neural real estate is proposed to represent the structure inherent across a wide range of knowledge domains including but not limited to spatial navigation (e.g., O’Keefe & Nadel, 1978; Epstein et al., 2017), social cognition (e.g., Park et al, 2021; FeldmanHall et al., 2025), and semantic cognition (e.g., Theves et al., 2019). Cognitive graphs represent knowledge in graph-like structures, in which item relations are captured by paths rather than by Euclidean distance (Peer et al., 2021). Neuroimaging work suggests that abstract neural representations of graph-like structures emerge in response to structured learning environments (Luettgau et al, 2024; Mark et al., 2025; Baram et al., 2025). Certain theories claim that humans use a repertoire of abstract structural forms to learn across a variety of knowledge domains (Kemp & Tenenbaum, 2008; Tenenbaum et al., 2011).

One can test for these putative abstract representations empirically: if, to some degree, structure is abstracted away from superficial features during or after learning, this will support the learning of new material that shares that same structure, even if all superficial features have changed (e.g., Mark et al., 2020; Collins & Frank, 2016; Garber & Fiser, 2025; Luettgau et al., 2024). These experimental designs are called “transfer paradigms”, because they test whether structure knowledge transfers to a novel environment. Using a visual statistical learning paradigm, Garber & Fiser (2025) found that implicit learners only demonstrated transfer of simple pair-wise associations after an intervening night of sleep. However, to our knowledge, only one prior study has used such a design to test transfer of higher-level graph structure knowledge in humans (Mark et al., 2020). In this study, participants were exposed to visual environments defined by two possible graph structures: a graph with community structure and a hexagonal lattice graph. Participants were better able to learn the second environment ∼24 hours later if its structure was congruent with the first environment, despite the fact that the visual items were nonoverlapping. The theoretical implication is that structural representations can be disentangled from surface-level information, and then applied in the service of new learning. The study also suggested that this process may require an intervening period of sleep, though this was not directly tested.

Despite evidence from other domains that structure knowledge can be abstracted, it remains unclear whether *semantic* structure is inextricably bound to or disentangled from its original features. One possibility is that learners represent the structure of a category by directly encoding relationships between pairs of specific features (e.g., increasing the association strength between “red” and “dots” after exposure to ladybugs). If semantic structure is encoded in this way, it is bound to specific features and the categories that contain those features; when those features or categories are not activated, the structure representation will not be activated either. Another possibility is that semantic structure becomes disentangled from specific features, either during or after learning, resulting in a more abstracted structural representation. In this case, structure representations could transfer to new learning contexts and support the acquisition of a new category, even if the new category is defined by a completely different set of features.

In the current experiments, we use a transfer paradigm to look for an abstracted representation of semantic structure and ask under what conditions it may arise. In particular, we explore whether sleep promotes the emergence of abstract structural representations and the extent to which memory reactivation during sleep in particular plays a role.

## RESULTS: Experiment 1

Experiment 1 tested for immediate transfer of structure knowledge. First, participants were exposed to either a non-modular “lattice” category (incongruent condition) or modular category (congruent condition) in a missing feature task (Fig. 2). Next, all participants were exposed to a new modular category in the missing feature task, which we will refer to as the transfer category. Structure knowledge of the transfer category was tested in a feature selection task and a feature arrangement task. If category structure is abstracted during or immediately after learning, then participants in the congruent group should exhibit increased performance on the transfer category relative to participants in the incongruent group, by virtue of having been previously exposed to a category defined by the same modular structure. In all regression analyses, the incongruent group was treated as the baseline.

**Fig. 2:**
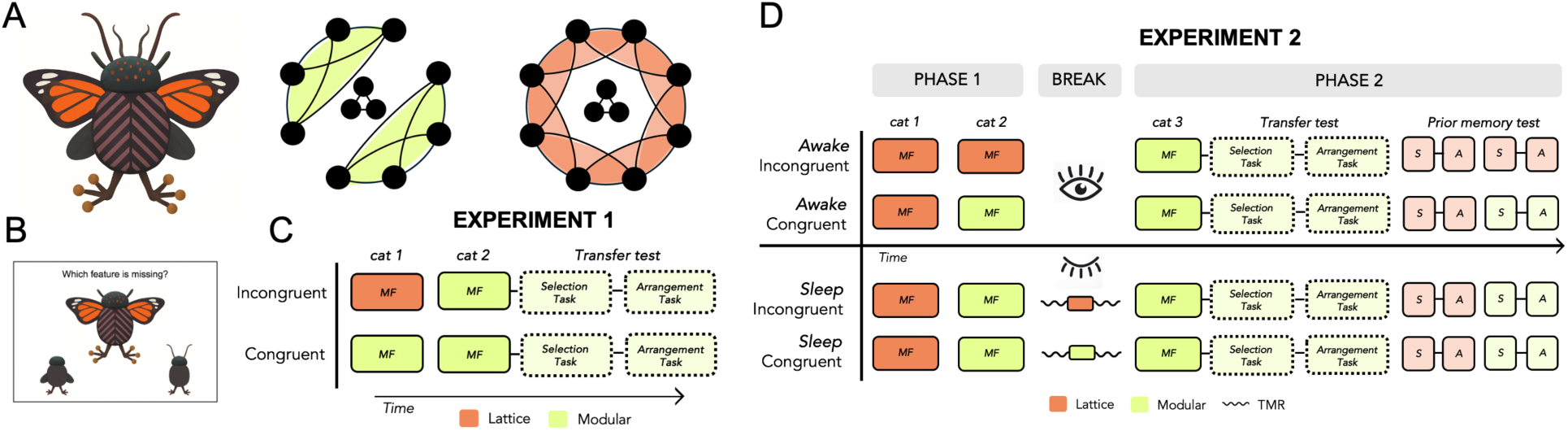
Novel categories and experimental design. In both experiments, humans learned the structure of novel categories in a missing feature (*MF*) task, and structure was tested in a feature selection (*S*) task and feature arrangement (*A*) task. (A) A cartoon example of one of the three novel categories used across the two experiments. The structure of each category was modular (green) or lattice (orange). Circles indicate features; lines indicate feature co-occurrence. Connections between the three central features and the peripheral features are balanced across structures and are not shown for visualization purposes. (B) Categories were learned in a missing feature task. (C) In Experiment 1, participants first learned either a lattice (incongruent) or modular (congruent) category, and then all participants learned and were tested on the transfer modular category with no delay. Boxes in dashed lines indicate the data relevant for the main transfer analysis; solid lines between boxes indicate tasks corresponding to the same category. (D) In Experiment 2, participants learned two categories in Phase 1 and were given a 3-hour break (wake or sleep) before learning the transfer category in Phase 2. During the nap, participants in the sleep-incongruent and sleep-congruent groups were played TMR cues corresponding to the previous lattice category and modular category, respectively.

Each participant’s responses on the feature arrangement task (i.e., Euclidean distances between features) were transformed into an 11 x 11 matrix capturing the relationships between the transfer category’s features. These response matrices were compared to the true structure matrix on both a group level and across participants. On a group level, a bootstrap analysis revealed that both the incongruent and congruent group response matrices captured the modular structure of the transfer category, compared to a permuted null baseline (Fig. 3A). However, neither group revealed significant alignment with the true structure across participants, and the congruent group did not outperform the incongruent baseline in a linear regression model (*t*(65)=0.44, *p*>0.6; Fig. 3B). No group difference on the feature arrangement task emerged when learning performance on the first category was added to the model (*p*>0.7).

**Fig 3:**
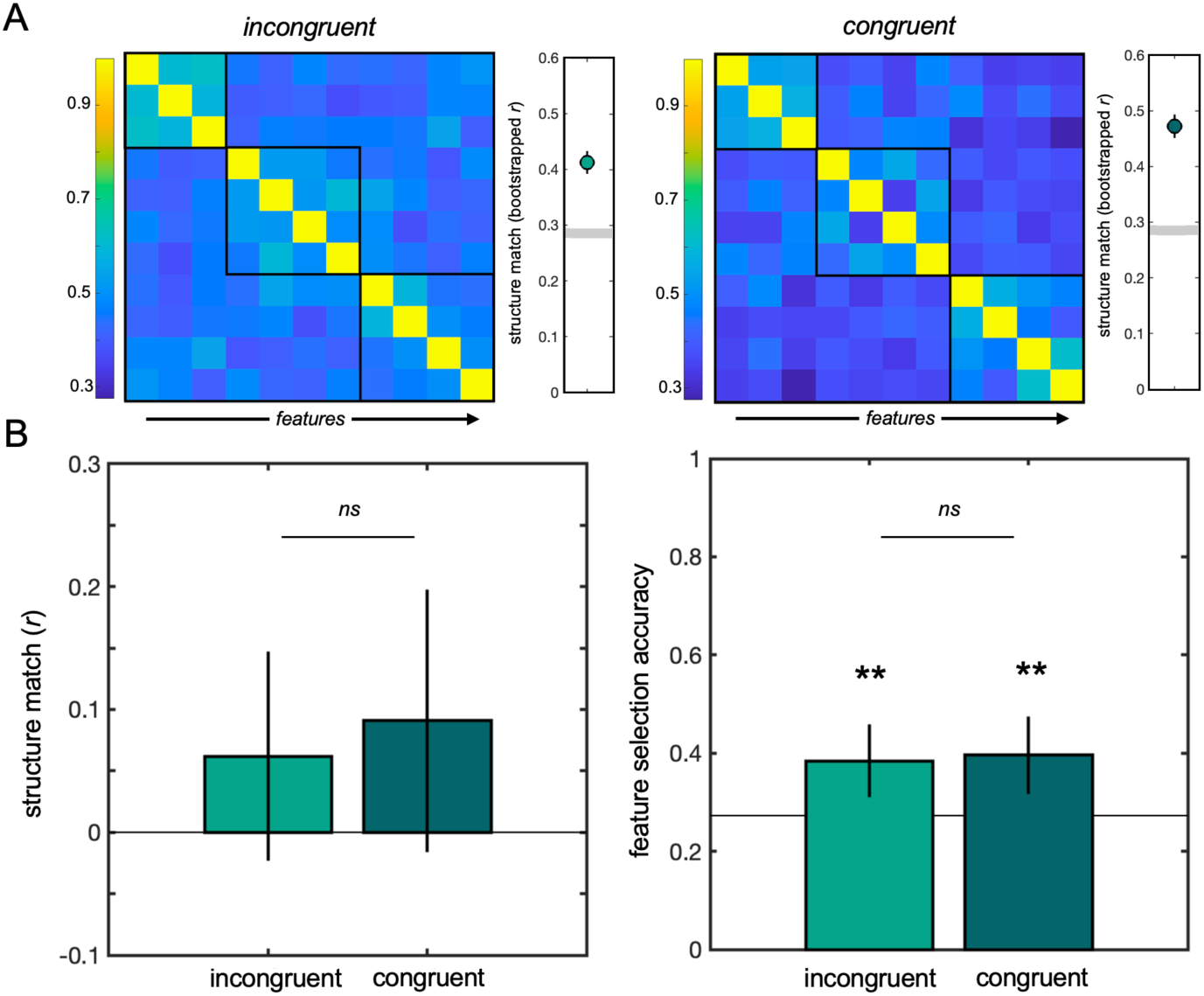
Experiment 1 results. No differences between the incongruent and congruent conditions were observed on the transfer category. (A) The average feature matrix derived from responses on the feature arrangement task matched the true modular structure of the transfer category within each condition. Lighter blue squares indicate stronger associations between features; darker blue square indicate weaker relationships. Black boxes indicate the true clusters of features that were associated in the transfer category. Datapoints to the right indicate significant structure knowledge in both conditions; grey lines indicate the significance threshold. (B) Neither group exhibited above-chance structure match across participants, and no difference between groups was observed. (C) Both groups performed above chance in the feature selection task, but no difference between groups was observed.

The two groups had comparable performance on the feature selection task: both the incongruent group (*t*(58)=3.03, *p*=0.004) and congruent group (*t*(47)=3.13, *p*=0.003) were able to identify the transfer category’s core features, and a logistic regression model revealed that the difference between the groups was not significant (*t*(105)=0.21, *p*>0.8). No group difference emerged on the feature selection task when learning performance on the first category was added to the model.

Overall, participants in the incongruent and congruent groups in Experiment 1 did not differ in their ability to learn the transfer category, thus providing no evidence of immediate transfer of structural knowledge. These results suggest that, immediately after learning, representations of structure are still tied to the surface-level features that were present in the learning environment.

## RESULTS: Experiment 2

Experiment 2 tested for structure knowledge transfer after a period of awake rest or after a nap. All participants learned two categories in Phase 1, and after a three-hour break learned the modular transfer category in Phase 2. Participants who remained awake during the break either learned two lattice categories (awake-incongruent group) or one lattice and one modular category (awake-congruent group) in Phase 1. This enabled us to test the hypothesis that structure representations transform with a delay. Participants who were given a nap opportunity during the break all learned one lattice and one modular category in Phase 1. Each category was paired with an auditory cue during learning; cues were ∼1 second clips of real insect sounds and occurred on 33% of Phase 1 trials. These cues were subsequently used in a targeted memory reactivation (TMR) paradigm in order to reactivate memories of a particular category during sleep. Some sleep participants were exposed to cues paired with the previously learned lattice category during their nap (sleep-incongruent group) and other participants were exposed to cues paired with the previously learned modular category during sleep (sleep-congruent group). Thus, the memories that were reactivated during sleep were either incongruent or congruent with the structure of the transfer category to which participants were exposed after their nap. In all regression analyses for Experiment 2, the incongruent group was treated as the baseline.

### Behavioral Results

The four conditions did not differ from each other in overall Phase 1 learning in the missing feature task (*F*(3, 84)=1.28, *p*>0.2), establishing equivalence between groups before the break. The order in which lattice and modular categories were learned in Phase 1 did not significantly affect Phase 2 transfer performance in the feature arrangement (*F*(1, 64)=3.43, *p*=0.07) or feature selection (𝜒^2^(1, 64)=1.68, *p*=0.20) tasks. The main question of interest was whether the baseline condition (awake-incongruent) was outperformed by any other condition (awake-congruent, sleep-incongruent, sleep-congruent) in learning the transfer category in Phase 2. Memory for the Phase 1 categories was also examined to further probe the effects of TMR.

### Transfer learning

We used regression analyses to ask whether experimental condition predicted transfer learning performance. In the feature arrangement task, a priori comparisons revealed that only the sleep-congruent group outperformed the baseline condition in terms of overall structure learning of the transfer category (*b*=0.14, *t*=2.02, *p*=0.046). Neither of the awake-congruent (*p*>0.2) or sleep-incongruent (*p*>0.3) groups differed from baseline. The full model did not reach significance (*F*(3, 84)=1.37, *p*>0.2). Examining each condition separately, only the sleep-congruent group exhibited successful structure matching, with correlations between participants’ structure and the true modular structure significantly greater than zero (*t*(22)=2.32, *p*=0.029; Fig 4B). Neither the awake-incongruent (*t*(23)=0.39, *p*>0.6), awake-congruent (*t*(20)=1.75, *p*=0.095), nor sleep-incongruent (*t*(19)=1.49, *p*=0.15) groups successfully captured the structure of the transfer category. Additionally, on a group level, a bootstrap analysis revealed that only the sleep-congruent group captured the modular structure of the transfer category, compared to a permuted null baseline (Fig 4A).

**Fig 4:**
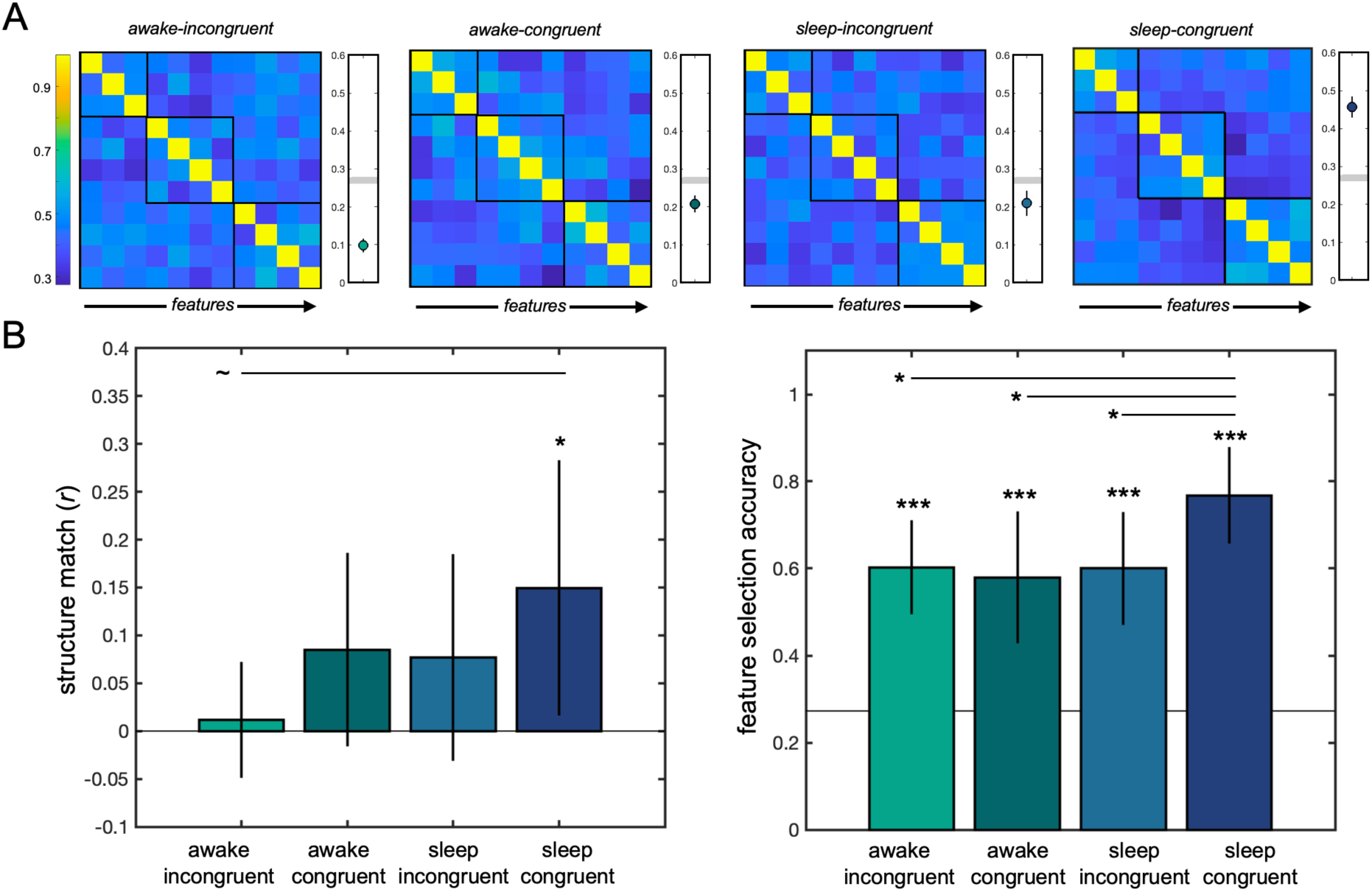
Experiment 2 behavioral results. The sleep-congruent group exhibited increased transfer learning. (A) Feature relationships extracted from the feature arrangement task were averaged across participants within each condition. Black boxes indicate the true clusters of features that were associated in the transfer category. A bootstrap analysis revealed that, on a group level, only the sleep-congruent group captured the transfer category’s modular structure. The grey line indicates the significance threshold; data points beneath that line indicate a lack of sensitivity to category structure. (B) In the feature arrangement task (left) the only group to successfully capture the overall structure was the sleep-congruent group. The difference between the sleep-congruent group and the awake-incongruent (baseline) group was marginal. In the feature selection task (right), all groups performed above chance, and the sleep-congruent group performed better than the awake-incongruent, awake-congruent, and sleep-incongruent groups. (* indicates p>0.05; ∼ indicates p< 0.1)

In the feature selection task, all groups performed at above chance levels (*p*’s < 0.001). The full regression model did not explain unique variance in transfer performance (𝜒^2^(3, 88)=7.2, *p*=0.07), but a priori comparisons revealed that the only condition that resulted in increased transfer performance, relative to the baseline condition, was the sleep-congruent group (*b*=0.78, *p*=0.033). Neither the awake-congruent group (*b*=-0.094, *p*>0.7) nor the sleep-incongruent group (*b*=-0.011, *p*>0.9) differed from baseline. Pairwise comparisons revealed that the sleep-congruent group’s accuracy (77%) was significantly greater than the accuracy of every other group: awake-incongruent (60%; *t*(47)=2.21, *p*=0.031), awake-congruent (58%; *t*(44)=2.08, *p*=0.043), and sleep-incongruent (60%; *t*(41)=2.07, *p*=0.045). Overall, behavioral results suggest that only the sleep-congruent group exhibited structure transfer from Phase 1 to Phase 2.

### Phase 1 memory

After participants completed the transfer learning tasks, memory for the two Phase 1 categories was tested. Because the Phase 1 memory test was administered after the transfer category tasks were completed, it is possible that the transfer category interfered with the two categories from Phase 1. However, these data provide an additional way to explore the behavioral effects of TMR (Supp. Fig 2). The awake-incongruent (baseline) group learned two lattice categories and no modular categories in Phase 1, thus performance on the two lattice categories were averaged, and this group was not tested on modular memory performance. Examining Phase 1 lattice memory, all groups performed above chance on the feature selection task (p’s < 0.001), and no comparison group differed from baseline (*p*’s > 0.3). No group exhibited overall structure match on the feature arrangement task for the lattice category assessed across participants (p’s > 0.3), including the sleep-incongruent group for whom the lattice category was cued during the nap (*t*(19)=0.034, *p*=0.97), and none of the comparison groups differed from baseline (*p*’s >0.4). Examining Phase 1 modular memory, above chance performance was observed in the feature selection task in the awake-congruent (*t*(22)=3.56, *p*=0.002), sleep-incongruent (*t*(19)=5.14, *p*<.0001), and sleep-congruent (*t*(22)=3.84, *p*<0.001) groups, and no difference between groups emerged (𝜒^2^(2, 63)=1.32, *p*=0.52). In the feature arrangement task, the only group to exhibit overall structure match to the Phase 1 modular category was the sleep-congruent group (*t*(22)=2.88, *p*=0.009). Responses in the awake-congruent (t(20)=0.72, *p*=0.48) and sleep-incongruent (*t*(19)=0.30, *p*=0.77) groups did not suggest that the structure was remembered. No difference between groups was observed (*F*(2, 63)=1.7, *p*=0.19). These results suggest that cueing the Phase 1 modular category during sleep in the sleep-congruent group boosted memory for that category upon waking, in addition to facilitating transfer to the new category.

### Effects of Sleep and Number of TMR Cues

#### Overall sleep time and stage correlations

The total amount of time spent asleep, and time spent in specific sleep stages, was determined for each participant (Supp. Fig. 1). Total amount of time spent asleep predicted memory for Phase 1 categories overall, across TMR cueing conditions (*b*=0.002, *p*=0.048). Examining the contribution of different sleep stages did not reveal a particular contribution from percentage of time spent in N2, N3, or REM (*p*’s>0.3). The total amount of time spent asleep marginally predicted performance on the transfer category in the feature selection task in Phase 2 (𝜒^2^(41, 2) =3.03, *p*=0.082). There were no specific contributions of percentage of time spent in N2 (*b*=0.38, *p*=0.79), N3 (*b*=01.77, *p*=0.39), or REM (*b*=3.4, *p*=0.11). No reliable relationships with sleep time were observed for the feature arrangement task.

#### Impacts of TMR Cueing

Next, we assessed the impact of TMR cueing on neural activity. The presentation of auditory cues significantly altered neural activity relative to sham: Target cues led to increases in power in the delta/theta band (cluster 1: 4-9 Hz, .48-.90 s after the cue, *p*=0.005) and in the slow spindle band (10-13 Hz, 1.59-1.91 s after the cue, *p*=0.03; Fig. 5A). No significant differences were observed between the response to target versus control cues. These increases in theta and spindle band power replicate prior work in the TMR literature and suggest that the sound cues successfully evoked neural responses (Siefert et al., 2024; Cairney et al., 2018; Ngo & Staresina, 2022). Since cues were predominantly administered in stages N2 and N3, we focus on these stages in the following analyses.

**Fig 5:**
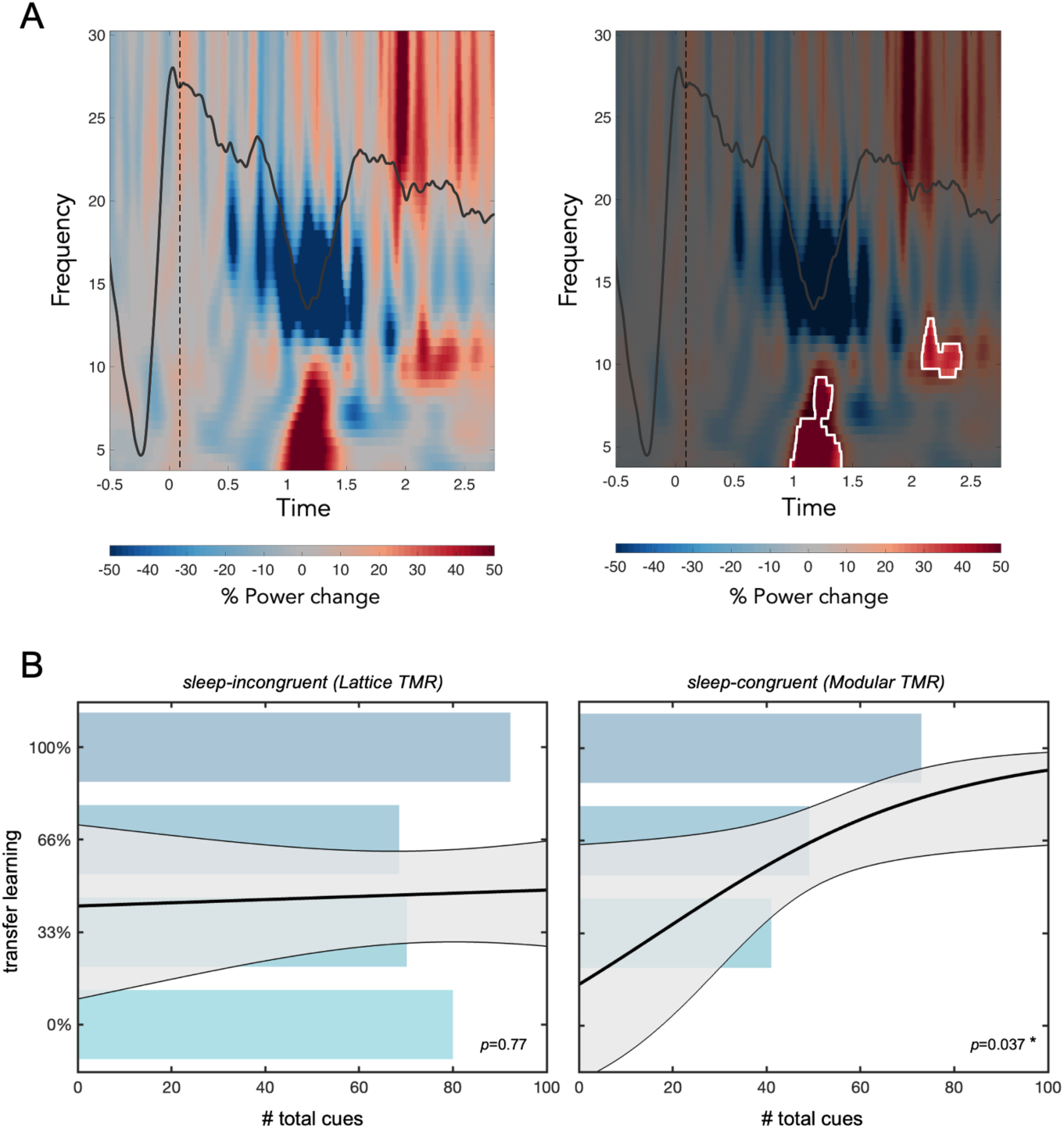
Effects of memory reactivation on transfer learning performance. (A) TMR cues produced a significant EEG response, relative to sham. The white lines in the right plot indicate significant clusters of increased power. (B) *Left:* In the sleep-incongruent condition, total number of TMR cues did not predict transfer learning performance in the feature selection task, controlling for total sleep time. *Right:* In the sleep-congruent condition, more cues predicted increased transfer learning performance. Bars are for visualization only and indicate the average number of cues presented to participants who performed at different accuracy levels on the feature selection task (no participants in the sleep-congruent group had 0% accuracy). The solid line and shaded region indicate the logistic regression model’s predicted accuracy and error at different numbers of cues.

**Fig 6:**
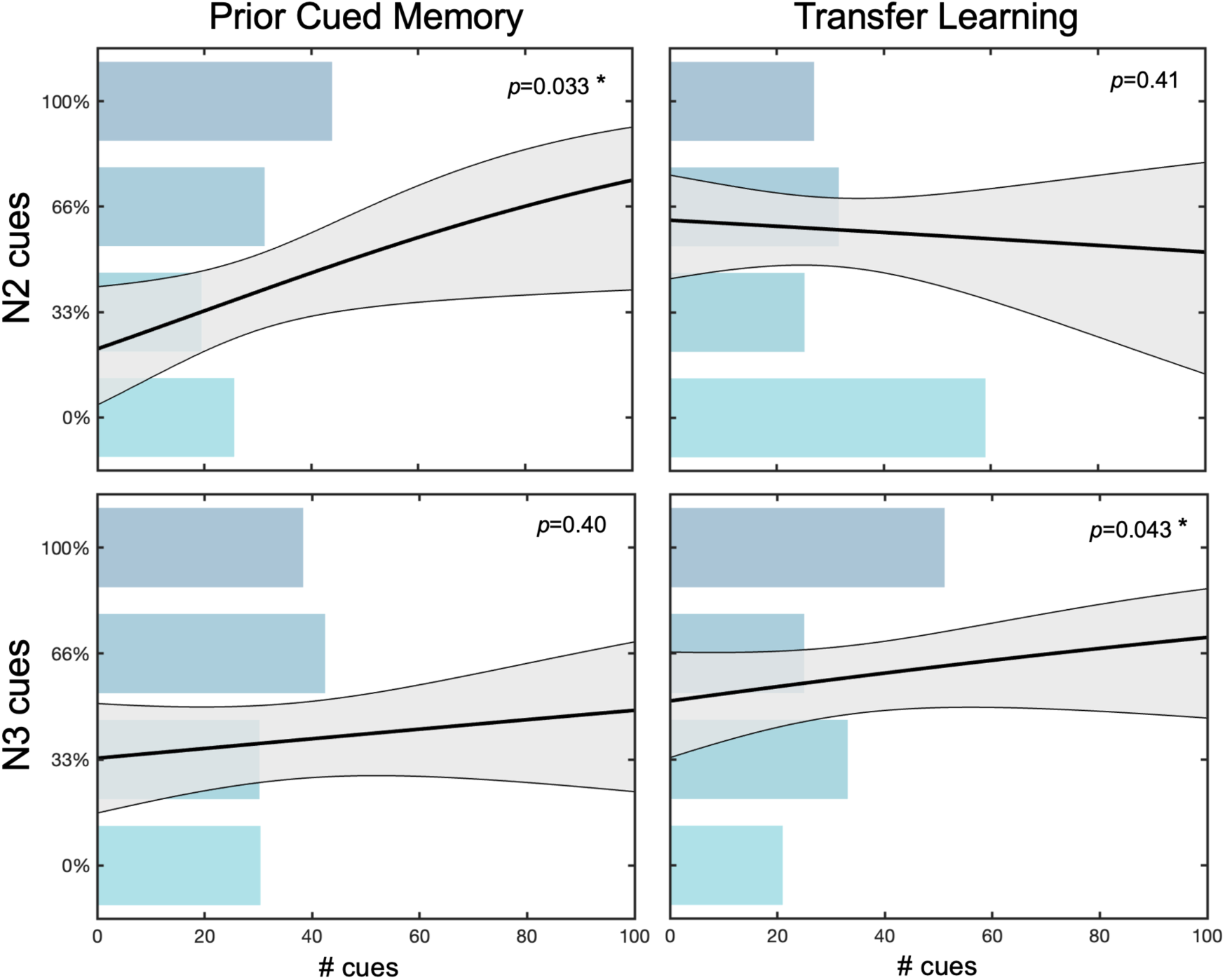
Effects of memory reactivation in different sleep stages. Cuing in N2 boosted Phase 1 memory, whereas cuing in N3 sleep facilitated transfer learning, as assessed in the feature selection task across all sleep participants. Plots on the left reveal the relationship between number of TMR cues and memory for the specific category from Phase 1 that was cued, split into cues presented during N2 sleep (top) and N3 sleep (bottom). Plots on the right reveal the relationship between number of transfer TMR cues and performance on the transfer category in Phase 2, split into cues presented during N2 sleep (top) and N3 sleep (bottom). In all plots, bars indicate the average number of N2/N3 cues presented to participants who performed at different accuracy levels on the feature selection task. The solid line indicates the logistic regression model’s predicted accuracy at different number of cues and the shaded region indicates the model’s predicted error.

Given the observed behavioral benefit of TMR on transfer learning in the sleep-congruent group, we asked whether the number of cues and the sleep stage in which the cues were presented impacted learning and memory. In particular, we asked whether reactivating memories of a learned category boosted memory for that specific category (i.e., Phase 1 memory), whether reactivating memories of a category with a certain structure facilitated learning for a new category with the same structure and no overlapping features (i.e., transfer learning), and how different sleep stages might differentially contribute to these effects. Across both sleep groups, target cues were always associated with Phase 1 memory (“Phase 1 cues”). For the sleep-congruent group, specifically, target cues were also congruent with the transfer category (“transfer cues”), in that the cued category from Phase 1 shared the same structure as the transfer category in Phase 2. Participants in the sleep-incongruent group were thus not exposed to any transfer cues.

##### TMR and Phase 1 Memory

Across both sleep groups, the total time spent asleep predicted memory in the feature selection task for the cued category (𝜒^2^(41, 2) =7.62, *p*=0.006) but not the un-cued category (𝜒^2^(41,2)=0.37, *p*=0.54) in Phase 1. These results are consistent with an overall effect of TMR on Phase 1 memory. When controlling for total sleep time, the total number of target cues presented was not predictive of Phase 1 memory (*b*=0.008, *p*=0.14). However, examining the impact of cuing in N2 and N3 sleep revealed that increasing the number of N2 cues boosted memory for the cued category (*b*=0.018, *p*=0.033), whereas the number of N3 cues did not affect Phase 1 memory in the same model (*b*=0.005, *p*>0.3), controlling for total sleep time. This suggests that memory reactivation during N2 sleep in particular may boost consolidation for surface-level features of that memory. These findings are consistent with many findings demonstrating that TMR successfully boosts memory for the cued content (Oudiette & Paller, 2013; Hu et al., 2020). No reliable effects of number of TMR cues on Phase 1 memory were observed in the feature arrangement task.

##### TMR and Transfer Learning

One of the unique questions that the current experiment can ask is whether the frequency or timing of TMR cues during sleep can facilitate future transfer learning. As mentioned above, across the two sleep groups, the total time spent asleep only marginally predicted transfer learning in the feature selection task (*p*=0.08). However, the total number of transfer cues did significantly predict transfer learning to a new category defined by the same structure but with a completely non-overlapping set of features (*b*=0.017, *p*=0.011), controlling for total time spent asleep. The full model explained a significant amount of variance in transfer performance (𝜒^2^(41,2)=10.6, *p*=0.005). Examining the sleep conditions separately, total number of cues predicted transfer performance for the sleep-congruent group (*b*=0.033, *p*=0.037) but not the sleep-incongruent group (*p*>0.7), controlling for total time asleep (Fig 5B).

Across the two sleep groups, the number of transfer cues presented during N3, specifically, predicted transfer learning performance, controlling for number of cues presented during N2 and the total time spent asleep (*b*=0.021, *p*=0.043). The number of transfer cues presented during N2 did not predict transfer learning in the same model (*b*=0.011, *p*>0.4). This differential contribution of cuing in N2 vs. N3 cannot be explained by time spent in these sleep stages: neither time spent in N2 (*b*=-0.006, *p*>0.4) nor in N3 (*b*=0.004, *p*>0.7) predicts transfer learning performance, controlling for total time asleep. Taken together, these results suggest that the number of memory reactivation events positively predicts transfer learning performance when the cued content and transfer content share a structural form. Memory reactivation during N3 sleep, in particular, appears to boost performance on a post-sleep transfer learning task. No reliable effects of number of TMR cues on transfer learning were observed in the feature arrangement task. Finally, an exploratory analysis probing the relationship between Phase 1 cued memory and transfer learning performance hints at a possible modulatory effect of sleep time (Supp. Fig. 2).

## DISCUSSION

We have shown, using a transfer learning paradigm, that semantic structure can be transferred across learning episodes, suggesting that this structure is abstracted and transformed away from surface-level features. This structure abstraction process does not happen immediately or automatically but is facilitated by memory reactivation during sleep.

In Experiment 1, participants learned two categories in immediate succession, and these two categories either differed in their feature-based structure (i.e., incongruent) or shared the same modular structure (i.e., congruent). If semantic structure is immediately abstracted during learning, participants in the congruent group would be expected to reveal better learning of the second, transfer category than participants in the incongruent group. However, this was not the case: participants in both the incongruent and congruent groups revealed successful structure learning of the transfer category, with no difference between the groups. These results suggest that, immediately after learning, representations of structure are still tied to the surface-level features of the learned categories.

Experiment 2 tested whether subsequent memory reactivation during sleep would facilitate abstraction. Participants in the baseline awake-incongruent condition learned only non-modular categories in the first learning phase and remained awake for three hours before learning the modular transfer category. Three other conditions were compared to the baseline in order to test whether structure is abstracted during a period of awake rest (i.e., awake-congruent), during a period of sleep (i.e., sleep-incongruent), or during a period of sleep in which memories of the Phase 1 modular category are reactivated (i.e., sleep-congruent). The only group that differed from baseline was the sleep-congruent group, suggesting that the reactivation of a recently learned category during sleep facilitates the abstraction of category structure. The more transfer cues presented, the better the transfer learning performance, with evidence that reactivation in N3 sleep in particular was helpful.

Experiment 1 found that abstract structural representations do not emerge immediately after learning, and the awake-congruent group in Experiment 2 revealed that these representations still do not emerge after a wakeful three-hour delay. Wakeful rest has been found to benefit memory consolidation (e.g., Brokaw et al., 2016; Dewar et al., 2012; Schapiro et al., 2018) and insight (Craig et al., 2018), particularly when the rest period is task-free rather than filled with other activities (Wamsley, 2019). Our findings suggest that awake rest periods are not sufficient for semantic structural transfer, though it is possible that less active wakeful breaks might have encouraged more transfer in our design.

Participants in the sleep-incongruent group in Experiment 2 were exposed to a modular category and a non-modular category in the first learning phase and then took a nap during which the previously learned non-modular category was reactivated using TMR. If general sleep consolidation processes facilitate the abstraction of structural knowledge, independent of memory reactivation, this group should reveal improved transfer learning performance of the modular category, relative to baseline—but this was not the case. Thus, sleep alone does not appear to facilitate the abstraction of structural representations, relative to wakeful rest, though it is possible that the incongruent cues interfered with an endogenous sleep-dependent abstraction process (TMR has been found to change the content of replay without changing the overall amount of replay, suggesting that reactivating one memory may lead to less endogenous reactivation of competing memories; Bendor & Wilson, 2012).

Participants in the sleep-congruent group in Experiment 2 were exposed to a non-modular and modular category in the first learning phase and then took a nap during which the previously learned modular category was reactivated using TMR. If sleep benefits structure abstraction via a mechanism of memory reactivation, participants in this group should reveal improved transfer learning relative to baseline. Indeed, the sleep-congruent group was the only group to successfully learn the overall structure of the transfer category in the feature arrangement task, and outperformed baseline and all other groups in demonstrating knowledge of the transfer category in the feature selection task (Fig. 4). Thus, while general sleep-based mechanisms did not facilitate transfer learning, reactivating the memory of a learned category during sleep appears to have promoted the abstraction of that category’s structure.

The TMR literature has revealed consistent benefits for memory for previously learned cued items (Hu et al, 2020). While the impact of TMR on Phase 1 memory was not our main question in the current study, our results do suggest that cuing a particular category facilitated later memory for that category. While there was no main effect of cuing on Phase 1 memory behavior, only the sleep-congruent group demonstrated memory for the Phase 1 modular category after the nap. Further, across all sleep participants, total sleep time predicted memory for the cued category, but not for the un-cued category, suggesting that reactivating a particular category with TMR prioritized that category for consolidation. We found that cuing in N2 sleep, in particular, facilitated this memory for the cued category.

Recent studies reveal that memory reactivation, via TMR, has impacts beyond the strengthening of unitary memories, and might contribute to the qualitative transformations of memories discussed above. In a category learning paradigm, we recently reported that cuing participants during NREM sleep with the names of category exemplars does not impact memories holistically. Rather, features unique to the exemplar were strengthened and features shared by all category members were weakened (Siefert et al., 2024). Another study suggests that while category exemplars are at first indelibly bound to the temporal context in which they were learned, reactivating the memories of those categories with TMR serves to disentangle temporal context from the category representations (Sherman et al., 2025).

Importantly, these findings highlight a potential tension in the literature regarding whether reactivation preferentially benefits item-specific information or more abstract, shared structure. One possibility is that the nature of the reactivation cue itself biases which level of representation is targeted, such that exemplar-specific cues and category-level cues may preferentially reinstate individuating features and shared information, respectively (Witkowski et al., 2021; Tulving, 1982). Thus, our use of category-level cues in this study may have been an important driver of our observed structure transfer effects. Nevertheless, our current results fit in with this pattern of empirical work—memory reactivation during sleep facilitated the transformation of category representations, resulting in an abstracted structural representation that could subsequently be used to benefit future learning.

Our data suggest a specific contribution of N3 sleep (Slow Wave Sleep; SWS), in that more cues presented in N3 was associated with better transfer learning performance. These findings are consistent with other data suggesting SWS-specific contributions to the abstraction of learned representations. In a study of cross-modal transfer, Durrant et al. (2016) exposed participants to a structure embedded in an auditory sequence of tones, and after a 30 minute or 24-hour delay tested whether participants were able to detect the same structure embedded in a visual display. Only participants tested after a 24-hour delay performed above chance, and increased time spent in SWS during the intervening night’s sleep predicted increased cross-modal transfer. In another study, participants who spent more time in SWS, relative to REM, exhibited increased explicit insight into an abstract regularity that was learned pre-sleep (Yordanova et al., 2008). Taken together, these findings suggest that SWS might facilitate the disentangling of specific features from a more abstract structural form (but see Hennies et al., 2017 and Lewis et al., 2018). Several theoretical accounts propose that memory reactivation during SWS supports forms of memory abstraction, including schema formation and gist extraction (Landmann et al., 2014; Inostroza & Born, 2013; Lewis & Durrant, 2011; Singh et al., 2022). These accounts often invoke hippocampal-neocortical interactions during hippocampally-driven memory reactivation that promote the formation of new, abstracted representations in neocortex (Baram et al., 2025). Future work will be needed to test whether our observed behavioral effects may be a result of this kind of systems consolidation process.

In conclusion, here we provide the first evidence that, in addition to direct memory benefits, memory reactivation during sleep via TMR can boost transfer learning in a novel environment with non-overlapping features. These results are consistent with a growing literature showing that sleep not only strengthens memories but transforms them.

## METHODS: General

### Category Structure

Two graphs were used to define category structure: a modular graph and a non-modular lattice graph (see Fig. 2A). Both graphs contain 11 nodes, three of which are three high-frequency “core” features that are always present and co-occur across all exemplars. The eight “peripheral” features in the modular graph cluster into two communities, such that an exemplar can have features from one community, or the other, but never from both. The eight peripheral features in the lattice graph were connected in a ring-like structure that did not result in clusters, or communities, of features. Though their peripheral structures differed, the graphs were matched on other network characteristics (for details, see Solomon & Schapiro, 2024). Each graph corresponds with eight unique six-feature exemplars, each of which contains three core and three peripheral features.

### Category Features

Two “species” of novel animals were used in Experiment 1, each defined by 11 discrete visual features presented on a base. The general kind and location of features were matched across species (e.g., horns, antennae, wings, arms), but there was no overlap of specific features across species. Six additional features per category were used for the catch trials described below. For a full description of these stimuli see Solomon & Schapiro (2024). An additional species, with the same characteristics, was designed for Experiment 2, enabling the creation of three categories with non-overlapping features. For each participant, species were randomly assigned to an appropriate graph, and the 11 visual features were randomly assigned to the 11 graph nodes.

### Category Learning

#### Missing Feature Task

Participants were exposed to each category in a missing feature task. Before the task began, participants were told: “We will train you on the animal categories by showing you category members with one feature missing. On each trial, you will be given two features and you will choose the feature that you think is the missing one.” On each trial, an exemplar with five features was presented. Two additional features, each presented on a desaturated base, were provided as response options. Participants clicked on the feature to make their response; trials were self-paced with no time limit. After a feature was selected, participants were notified whether their response was correct or incorrect. If the incorrect feature was chosen, the trial was repeated. Each category was tested across 102 total trials, including 96 experimental trials (48 core; 48 peripheral) and six catch trials. In the core structure trials, the exemplar was missing one of its three core features, and the incorrect option was a structure-consistent peripheral feature. These trials trained participants to learn that all exemplars must have all three core features. In the peripheral structure trials, the exemplar was missing one of its peripheral features—the correct option was a structure-consistent peripheral feature, and the incorrect option was a structure-inconsistent peripheral feature. These trials trained participants to learn the modular or lattice peripheral structure of the assigned graph. Six additional catch trials were used to confirm that participants were following instructions and paying attention. In each catch trial, the incorrect option was a novel feature that had never been seen before in the experiment. Participants were instructed at the beginning of the task to not select these novel features when they appeared. The order of all trials was randomized, the only constraint being that catch trials only appeared in the second half of the task.

To assess category learning in the missing feature task, each trial was coded as accurate (1) or inaccurate (0) based on the participants’ initial response. For each participant, for each category, mean accuracies on experimental trials and catch trials were separately calculated. Participants who exhibited above threshold catch trial performance (>50%) for each category were included in the main analyses; participants who did not pass this threshold were excluded. Mean accuracies in each condition were compared to chance (50%) using t-tests, and performance across conditions was compared using a linear regression model.

### Testing Structure Knowledge

Category knowledge was tested in a feature selection task and a feature arrangement task. The feature selection task was a strong yet simple test of category structure, whereas the feature arrangement task provided a more fine-grained measure reflecting whether the overall structure of each category was learned and retained.

#### Feature Selection Task

In the feature selection task, the category’s 11 features were visually displayed in a grid, in a randomized order, and participants were asked to “click on the three features that are most important for this species.” When a feature was selected, a blue outline appeared around the feature so participants could keep track of their responses. Once three features were selected, the task immediately ended. Responses on the feature selection task were analyzed by, for each participant, recording the number of correctly selected core features (out of three) and calculating percent accuracy. Mean accuracies in each condition were compared to chance (27%) using t-tests, and performance across conditions was compared using a logistic regression model.

#### Feature Arrangement Task

In the feature arrangement task, the category’s 11 features were randomly displayed around the edges of a large white circle, and participants were asked to click and drag the features into the circle, putting related features together. They were told to base their decisions on what they learned during the experiment (https://meadows-research.com). A separate set of AMT participants (*N*=80) completed a version of this task in which the instructions were to arrange features based on visual similarity. This enabled us to control for visual dissimilarity when analyzing each participant’s task-based dissimilarity data.

Each participant’s feature arrangement data were analyzed by extracting the Euclidean distances between all pairs of features, and z-scoring these distances within subject, resulting in an 11 x 11 feature dissimilarity matrix. Participants’ data were compared with the true feature dissimilarity matrix, derived from the underlying graph structure. In a group-level analysis, dissimilarity matrices were averaged within each condition before quantifying the group’s alignment with the true structure. First, a significance threshold was determined based on a permuted null distribution of all participants’ dissimilarity scores. Feature labels were randomly permuted on the true dissimilarity matrix before correlating it with participants’ data across 1000 iterations; the significance threshold was set as the 95^th^ percentile of this permuted null distribution. The alignment between the group’s mean dissimilarity matrix and the true dissimilarity matrix was calculated using Pearson’s r, and a bootstrap analysis was used to generate confidence intervals, in which *N* individual participants’ data was sampled with replacement and averaged before calculating Pearson’s r across 1000 iterations, where *N* is the group’s sample size. In a participant-level analysis, we determined whether the true feature dissimilarity matrix could predict participants’ feature dissimilarity matrix, controlling for visual dissimilarity between features. A linear regression analysis was run for each participant, and the resulting beta corresponding to the true structure was extracted. Mean betas in each group were compared to zero using t-tests, and conditions were compared using a linear regression model.

### Statistical Analysis

Data from the missing feature, feature selection, and feature arrangement tasks were analyzed for each category for each participant. In our main analyses, regression models were used to determine the influence of experimental condition on task performance. A linear regression model was used to predict missing feature data and feature arrangement data, since these are continuous variables. The feature selection task data reflected a proportion correct out of three attempts and could thus take one of four values (0, 0.33, 0.66, 1). Because these proportions arise from a binomial process rather than a continuous distribution, it was modeled as a stochastic outcome and estimated via logistic regression. In all regression models, condition was added as a categorical predictor.

## METHODS: Experiment 1

### Participants

107 participants between 18-35 years of age contributed data to Experiment 1 and were recruited on Amazon Mechanical Turk (Mean age: M=25.5, SD=2.2, 62% female). An additional 104 participants were excluded because they either did not reach criterion on catch trial accuracy in the missing feature task, reported taking notes during the experiment, or reported not taking the experiment seriously. Out of the final set of 107 participants, 40 participants did not provide data on the feature arrangement task due to technical error. Participants received a base pay of $0.50 in addition to a bonus of up to $5.00 based on task completion and performance on the orthogonal attention check. Consent was obtained for all participants in accordance with the University of Pennsylvania IRB.

### Experimental Design

In Experiment 1, participants learned two categories in immediate succession. The structure of the first category was randomized (i.e., modular or lattice), but the second category, the “transfer” category, was defined by a modular structure for all participants. This resulted in two experimental conditions in which the first category’s structure was either congruent or incongruent with the transfer category: participants who first learned a lattice category were in the “incongruent” group, whereas participants who first learned a modular category were in the “congruent” group. Participants completed the missing feature task for the first category and then completed the missing feature task for the transfer category. Participants then completed the feature selection task followed by the feature arrangement task for the transfer category.

### Behavioral Analysis

For each participant we assessed category learning in the missing feature task for the first category in addition to the transfer category. The main analyses focused on structure knowledge of the transfer category, assessed via the feature selection task and feature arrangement task. In all regression analyses, the incongruent group was treated as the baseline. If learned structure representations are immediately disentangled from surface-level features after learning, participants in the congruent group, who learned two modular categories in succession, are expected to outperform participants in the incongruent group in terms of transfer category structure knowledge.

## METHODS: Experiment 2

### Participants

Ninety-two (92) participants from the University of Pennsylvanian community contributed data to the main analyses of Experiment 2 (Mean age: *M*=22.6, *SD*=4.3, 54% female). An additional 19 participants were excluded because they did not reach criterion on catch trial performance in the missing feature task. Across the two sleep conditions, additional participants were excluded if they did not sleep long enough for TMR cues to be presented (*N*=23) or if their reports indicated that they heard the TMR sound cues during the nap period (*N*=12). Out of the final set of 92 participants (awake-incongruent: *N*=26; awake-congruent: *N*=23; sleep-incongruent: *N*=20; sleep-congruent: *N*=23), 4 participants did not provide data on the feature arrangement task due to technical error. Participants received course credit or monetary compensation for participating in the experiment. Consent was obtained for all participants in accordance with the University of Pennsylvania IRB.

### Experimental Design and Procedure

The experiment consisted of one 5 h session that contained a pre-break learning phase (Phase 1), a 3 h rest period, and post-break learning and test (Phase 2). Two categories were learned in Phase 1, and one category was learned in Phase 2 (the “transfer” category). The structures of the Phase 1 categories and the nature of the rest period were manipulated across four conditions to test different hypotheses regarding structure transfer learning (i.e., awake-incongruent, awake-congruent, sleep-incongruent, sleep-congruent; Fig. 2D).

Participants arrived in the laboratory between 12:00pm-1:00 PM. Participants had been notified whether they had been randomly assigned to a wake or sleep condition prior to arrival. They were informed that the experiment related to learning and memory but were neither informed of their assigned experimental condition nor that TMR might be used. Consent was obtained before the Phase 1 learning tasks (∼1 h). After Phase 1 was complete, all participants were given a 3 h rest period. Participants in awake conditions spent this time occupying themselves with materials they had brought with them (e.g., homework, books). For participants in sleep conditions, ∼40 mins were spent on EEG capping before a 2 h nap opportunity was provided in a designated sleep room in the laboratory. After the nap, the cap was removed, and participants were given 20 mins to recover from sleep inertia. After the break, all participants completed survey asking about their experiences. In Phase 2, all participants learned the final transfer category, and structure knowledge of all three categories was tested.

#### Phase 1 Learning

All participants learned two categories in Phase 1 by completing 102 trials of the missing feature task for each in succession. Participants in the awake-incongruent condition learned two lattice categories in Phase 1, obviating the possibility of structure transfer to Phase 2. This condition thus served as our baseline. All other conditions (i.e., awake-congruent, sleep-incongruent, sleep-congruent) learned one lattice and one modular category in Phase 1 in a randomized order. Each category in Phase 1 was randomly paired with one of three possible auditory cues (see below). During the missing feature task, the designated cue had a 33% probability of playing on each trial.

#### Rest Period

All participants took a 3 h break in between Phase 1 and Phase 2. Participants in the two awake conditions were told they could either remain in the testing room or relocate to another nearby seating area. During the break, participants had access to books and screens. Participants in the two sleep conditions underwent EEG capping and were given a 2 h nap opportunity. During their nap, two distinct cues were presented following the TMR procedure described below. One of these cues was a cue previously paired with a category from Phase 1 (target cue) and the other cue was unfamiliar (control cue). For participants in the sleep-congruent group, the target cue was the cue previously paired with the modular category in Phase 1; for participants in the sleep-incongruent group, the target cue was the cue previously paired with the lattice category in Phase 1. In this way, the cued category was either incongruent or congruent with the structure of the transfer category. Sleep participants recovered from sleep inertia before beginning Phase 2. Participants who explicitly reported hearing the TMR cues, or reporting hearing a sound that could be identified as a TMR cue, were excluded from analysis (*N*=12).

#### Phase 2 Learning and Test

The design of Phase 2 was identical for all participants in all conditions. First, participants completed 102 trials of the missing feature task for the final, modular transfer category. No auditory cues were paired with this category. Immediately after learning, participants completed the feature selection task followed by the feature arrangement task for the transfer category. Once completed, participants were tested on their memory for the two Phase 1 categories in a randomized order. For each category, the feature selection task was completed first, followed by the feature arrangement task.

### EEG Recording

EEG was recorded during the rest period for participants in one of the two sleep conditions. Data were collected via an actiCHamp amplifier (Brain Vision) using a 64-channel EasyCap active electrode EEG cap with a sampling rate of 512 Hz and an online reference placed at FCz. Two electrodes were repurposed to record electro-oculography and two to record electromyography for the purposes of sleep scoring, and two electrodes were placed on the mastoid. EEG capping took place after Phase 1 learning at the beginning of the 3-hour rest period, and impedance was checked before the nap commenced and was kept below 20 kΩ.

### Auditory Cues

The sound cues paired with the categories in Phase 1 were insect sounds sourced from the Bug Bytes sound library (Mankin, 2019). The three sound files used were naturalistic recordings of an ant, a wasp, and a cricket. For each auditory file, a 1 second clip was selected such that the sounds were distinctive from each other yet still identifiable as insect sounds. The sound files were edited to be matched in volume. For each participant, the three sounds were randomized to one of three assignments: the Phase 1 category target cue, the Phase 1 category control cue, or the TMR control cue. Both of the categories in Phase 1 were paired with a cue, but only the target cue was replayed during the nap, alongside, the TMR control cue. That is, participants had not been exposed to the TMR control cue before the nap began.

### Real-time TMR

Endogenous memory reactivation during sleep occurs in the up-states of slow-oscillations (Diekelmann and Born, 2010; Mölle and Born, 2011; Gulati et al., 2014; Lewis and Bendor, 2019; Schreiner et al., 2021), and administering TMR cues in this oscillatory phase has been found to be especially beneficial for memory consolidation (Batterink et al., 2016; Göldi et al., 2019; Ngo and Staresina, 2022; Abdellahi et al., 2023; Xia et al., 2023). Spindles occur in N2 and N3 sleep and are thought to also be related to memory reactivation (Born and Wilhelm, 2012; Klinzing et al., 2019). Spindle activity is followed by a refractory period, during which TMR cues are not efficacious (∼2.5s; Antony et al., 2018b). We adopted the TMR protocol from Siefert et al. (2024) to play auditory cues both in the up-states of slow oscillations and outside of the spindle refractory period.

OpenViBE Brain Computer Interface Software (Renard et al., 2010) and custom MATLAB scripts were used to track slow oscillations and spindles in parallel. If the Fz-filtered signal crossed a threshold of +35 μV, an SO was detected, and if at least 2.5 s had passed since the last detected spindle, the system played an auditory cue at the SO peak (i.e., the time at which the signal’s slope changed from positive to negative). The TMR system waited for 8s after each sound before detecting a new SO. In addition to playing sounds (i.e., target and control cues), spaced throughout the cueing list were “no sound” triggers: at certain monitored SO up-states, no sound was played but the timepoint was marked. These “no sound” SOs were used as a sham for computing sound-evoked responses. This system was implemented using MATLAB 2011a (64 bit), OpenViBE Designer v2.2.0 (64 bit), and OpenViBE for Brain Products server (64 bit).

### Sleep EEG data analyses

Analyses of sleep EEG data were performed in MATLAB (2022b) using the FieldTrip toolbox (Oostenveld et al., 2011).

#### Sleep stages

For sleep scoring, data were downsampled to 256 Hz, re-referenced to the average of the linked mastoids, and filtered between 0.1 and 35 Hz. Sleep scoring was done using Dananalyzer (https://github.com/ddenis73/danalyzer; Denis et al., 2021) in 30-second epoch increments by expert scorers according to standard criteria (Iber et al., 2007). For each participant, the number of epochs spent in N1, N2, N3, and REM was recorded, and total sleep time was estimated as the total time spent in these four sleep stages. By scoring the sleep records, we could also determine how many target, control, and sham cues were played in the different sleep stages across participants.

#### Time-frequency analyses

For both time-frequency representations (TFRs) and event-related potentials (ERPs), data were cut from −3 to 3 s centered on the time of cue delivery. Data were then downsampled to 256 Hz, re-referenced to the average of the linked mastoids, and filtered between 0.3 and 30 Hz. Artifacts were rejected in two stages: first, the experimenter manually looked through every individual trial and channel and made note of those which looked problematic. Next, trials and channels were confirmed for removal via FieldTrip’s visual summary functions, using amplitude, variance, and kurtosis. For TFR analyses, data were convolved with complex Morelet wavelets to obtain spectral power from 4 to 30 Hz in 0.5 Hz frequency steps and 5 ms time steps. Then, participant-specific TFRs were converted into percent power change relative to a −1.25 to −.25 ms pre-cue window (Sherman et al., 2025). For analyses of sound-elicited activity, TFRs for target sounds, control sounds, and no sound (sham) trials were processed separately. Different TFRs were obtained for each participant (TFR target-sound minus TFR sham; TFR target-sound minus TFR control-sound). For the visualization of ERPs in Fig. 5, data were high-pass filtered at .5 Hz and baseline corrected with respect to the -200 to 0ms baseline before cue onset (Cairney et al., 2018; Siefert et al., 2024).

To identify significant time-frequency clusters of sound-elicited activity, trial-specific spectrograms were averaged across electrodes within participants. A t-test was then performed across subjects, generating candidate clusters of points in time-frequency space where power was different from zero (α = 0.01). To account for multiple comparisons, cluster size-based permutation testing was implemented: a permutation-corrected p-value was determined for each candidate cluster by calculating the frequency of the candidate cluster size exceeding maximum cluster sizes in a permutation distribution obtained by running one-sample t-tests after randomizing the sign of power values among participants (1,000 permutations; significance level: *p*=0.05). We examined sound elicited activity by analyzing the differences between target sounds and sham, as well as between target sounds and control sounds.

## Acknowledgments

We are grateful to Paige Sevchik for sleep scoring assistance. This work was supported by NIH grant R21 MH128788.

## SUPPLEMENTARY MATERIALS

**Supp. Table 1:** Time and cues in different sleep stages. Number and proportion of minutes spent in each sleep stage, and number and proportion of cues played in each sleep stage. Parentheses indicate the standard deviation.

|  | Total Sleep | N1 | N2 | N3 | REM |
| --- | --- | --- | --- | --- | --- |
| Mean minutes of sleep | 90.36<br>(23.11) | 14.72<br>(9.75) | 51.30<br>(20.38) | 13.09<br>(10.91) | 11.24<br>(11.12) |
| Mean proportion of sleep | -- | 0.18<br>(0.14) | 0.57<br>(0.17) | 0.14<br>(0.11) | 0.11<br>(0.11) |
| Mean number of cues | 66.21<br>(37.54) | 0.33<br>(2.0) | 29.14<br>(24.0) | 35.91<br>(35.31) | 0.84<br>(5.03) |
| Mean proportion of cues | -- | 0.01<br>(0.08) | 0.50<br>(0.30) | 0.47<br>(0.32) | 0.02<br>(0.13) |

**Supp. Figure 1:**
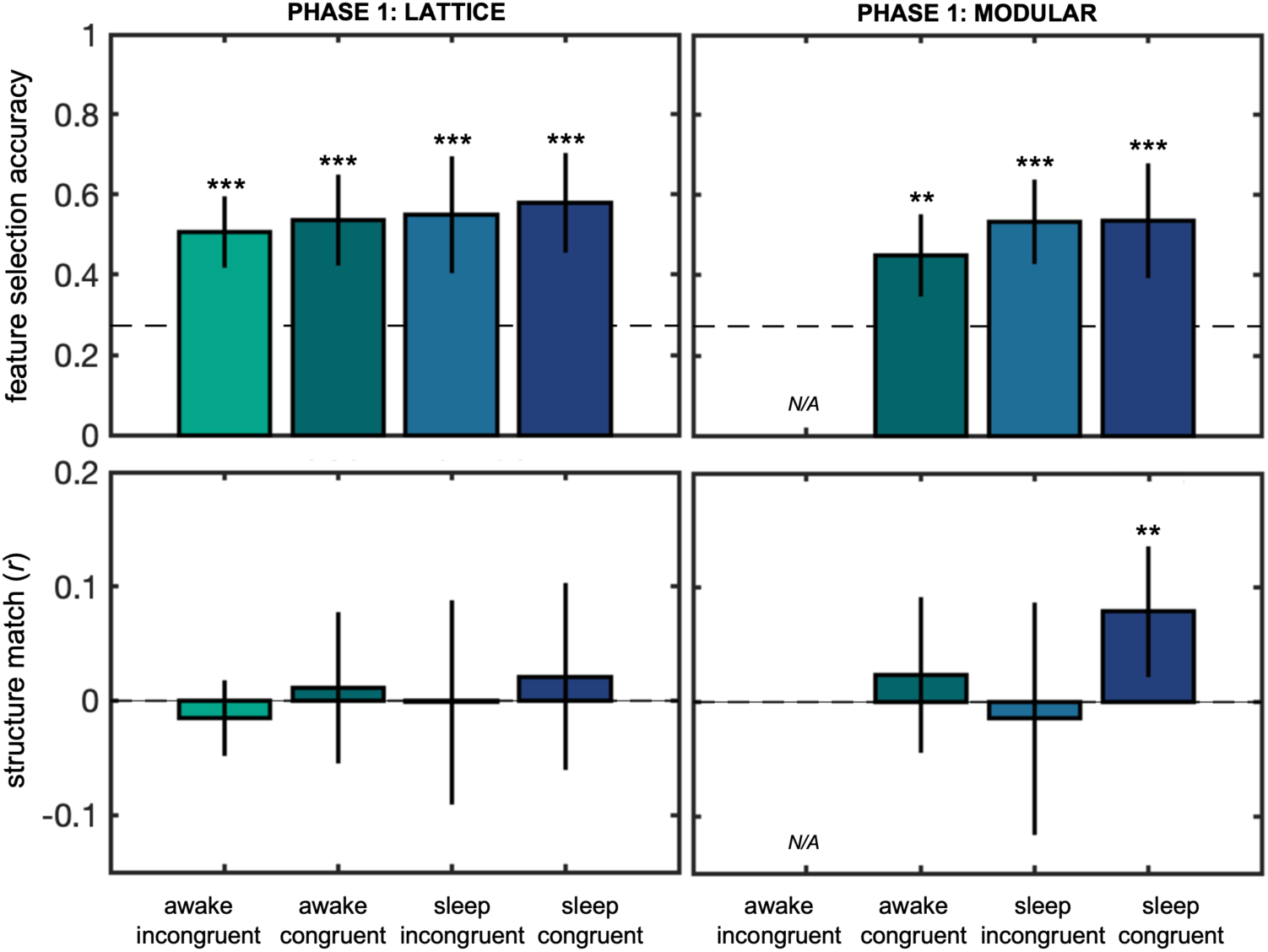
Memory for Phase 1 categories. Mean feature selection accuracy (top) and structure match data (bottom) for the Phase 1 Lattice (left) and Modular (right) categories. Participants in the awake incongruent condition did not learn a Modular category in Phase 1.

**Supp. Figure 2:**
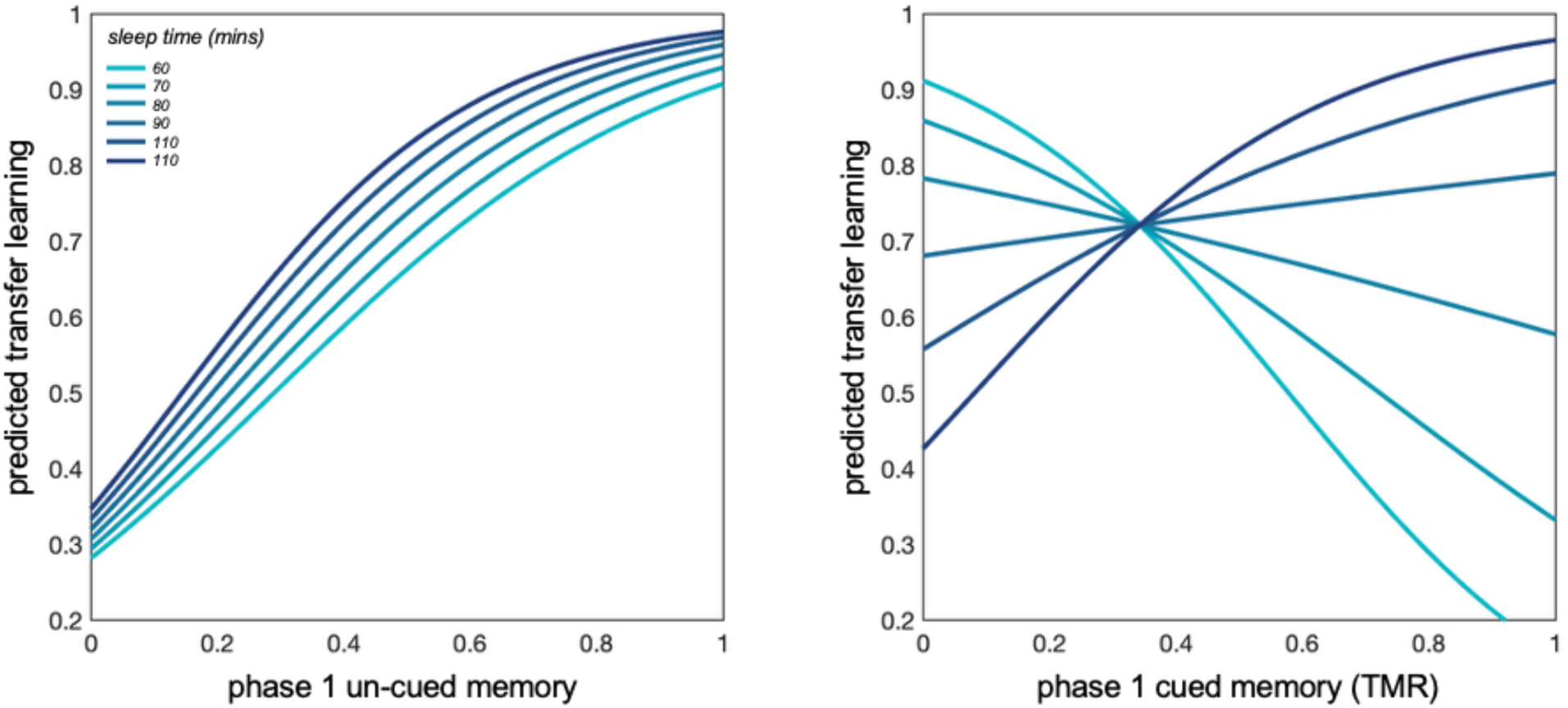
Sleep interacts with Phase 1 memory to predict transfer learning. Relationships between Phase 1 memory, transfer learning, and sleep were observed in the sleep-congruent group. *Left:* Memory for the un-cued category from Phase 1 was strongly linked to transfer learning performance in Phase 2, and this relationship was not modulated by sleep. *Right:* The correspondence between memory for the cued category in Phase 1 and transfer learning in Phase 2 is modulated by total sleep time. At shorter periods of sleep (∼80 mins or less), better cued memory predicts worse transfer learning; as sleep length increases (∼90 mins or more), better cued memory predicts better transfer learning performance. These results reveal the potential time course by which surface-level representations are transformed into more abstracted structural representations.

## Phase 1 Memory and Sleep Interact to Influence Transfer Learning

In the sleep-congruent group, any benefit of TMR on transfer learning is presumably due to reactivating the memory of the previously learned cued category. We would thus expect a positive relationship between memory for the cued category and transfer learning performance: if the target cues result in transfer, it is because the memory of the Phase 1 category was reactivated, thus this memory should be strengthened. On the other hand, we would expect a weaker relationship between transfer learning and the un-cued category, since the two are not directly causally related. Instead, we found only a marginal relationship between cued memory and transfer learning in the feature selection task (𝜒^2^(2, 21)=4.06, *p=*0.053), but a strong relationship between un-cued memory and transfer (𝜒^2^(2, 21)=11.3, *p=*0.001). Similar results were found when we controlled for total sleep time separately for the cued category (*b*=1.84, *p*=0.062) and the un-cued category (*b*=4.16, *p*=0.002). These puzzling results suggest a nuanced relationship between memory for the cued category and the transfer learning that this memory reactivation facilitates. To explore this further, we ran an exploratory post hoc analysis testing for a possible interaction with sleep time. The interaction between total sleep time and prior cued memory was significant: when total sleep time, prior cued memory, and their interaction were used to predict transfer learning performance in the feature selection task, there were marginal main effects of prior cued memory (*b*=-13.27, *p*=0.054) and total sleep time (*b*=-0.026, *p*=0.066), but the interaction between them was significant (*b*=0.077, *p*=0.027). The full model including the interaction term explained a significant amount of variance (𝜒^2^(2,21)=9.6, *p=*0.022) whereas the model excluding the interaction term did not (𝜒^2^(2,21)=4.08, *p=*0.13). The observed interaction indicates that the correspondence between prior cued memory and transfer learning depends on the amount of sleep (Supp. Fig 2B). When sleep is short (<80 mins), stronger prior cued memory predicts worse transfer learning; however, with longer sleep (>90 mins), the pattern reverses, and stronger prior cued memory predicts better transfer learning. On the other hand, the positive relationship between prior un-cued memory and transfer learning is not modulated by sleep (*b*=0.012, *p*>0.7; Supp Fig 2A). Given our previous finding that time spent in N2 or N3 did not impact transfer learning directly, these data potentially reveal the time course by which structural representations are transformed away from surface-level details and into a more abstracted form.

